# Sign and strength of pairwise interactions in natural isolates depend on environment type

**DOI:** 10.64898/2026.03.31.715556

**Authors:** Terence McAvoy, Elze Hesse, Angus Buckling, Luke Lear

**Author notes:** **Author Correspondence:** ESI Carpark, Mabe Burnthouse, Penryn TR10 9FE.

## Abstract

Bacterial interactions-whether positive or negative – are crucial for the functioning of microbial communities. Though bacterial interactions are mainly expected to be negative, the sign and strength of interactions are predicted to be context dependent, with interactions typically being more positive in more stressful and nutrient-poor conditions. However, systematic studies investigating how the environment affects interactions between multiple taxa are lacking.

Here, we determine if interactions between a panel of natural soil isolates change in response to the environment in which they are grown, with two different artificial media used (one simple and one complex) and a more ecologically relevant soil wash. To maximise natural variation in interactions, we collected multiple isolates from multiple sites: co-occurring (sympatric) isolates were predicted to show more negative interactions than allopatric isolates because of greater overlap in resource use. Pairwise interactions were in general negative, but more negative when grown in a complex lab-derived medium (Tryptic Soy Broth). Mutually beneficial interactions were most common in a simple resource medium (M9 minimal media) and exploitative interactions were most frequent in a soil broth. These patterns were independent of whether species originated from the same or a different site. The study supports the prediction that nutrient rich environments promote more negative interactions, and that measuring interactions of soil isolates in standard lab media is likely to misrepresent interactions occurring in natural environments.

## Introduction

Bacterial communities underpin many key ecological processes and live in diverse environments that vary in their resource complexity and availability. Interactions between co-occurring species can range from cooperative and mutually beneficial (+/+), through exploitative (+/-) to competitive (-/-). The prevailing interaction type can shape community composition and ecosystem services (1–3), and influence overall stability, with competitive interactions predicted to increase the latter (4, 5). Understanding interspecific bacterial interactions is therefore key to predicting and maintaining bacterial community function in both natural (6) and applied contexts (7). Though similar interactions can occur within-species, these primarily involve different mechanisms (e.g. kin-selection (8)) that are not the focus of the current study.

Resource characteristics, including complexity, availability, and overall abundance, are key factors influencing the type and strength of bacterial interactions (9, 10). For example, environments which are rich and diverse in nutrients will support high bacterial densities, which in turn increase the magnitude of competition for space and resources (11), leading to competitive interactions (-/-) (12–14). More complex nutrient-rich environments can allow bacterial investment both into growth and competitor suppression (e.g. through production of toxins), as observed in *Bacillus sp.*(*15*). Alternatively, environments containing resources which are less readily available can lead to bacterial species alleviating competition through niche complementarity, for instance as species break down recalcitrant resources, or as they rely on conspecifics for enzymatic breakdown. (9, 10).

Both mutually beneficial (+/+) and exploitative (+/-) interactions are equally dependent on environmental resources, and on the resulting flow of metabolites between species. Metabolic cross feeding, where species (co) consume metabolic byproducts of others (16), can lead to mutual benefits for interacting species, as in seen between methanogenic archaea and sulphate-reducing bacteria (17, 18), where by-products produced by the reducing bacteria are used by methanogens, preventing harmful waste buildup that would inhibit bacterial growth; whilst adaptation to utilise waste products has also been shown to decrease competitive interactions amongst bacterial isolates in beech tea medium (19). Such metabolic dependencies are sensitive to changes in the resource environment and interaction types can alter in response to changes in external nutrient supply (20, 21). For example, the mutualistic interaction between *E. coli* and *S. enterica* shifts to exploitative (22) when adding limiting metabolites: while *S. enterica* still benefits from the acetate produced by *E. coli,* the latter gains no return and competes for shared nutrients. A similar pattern was observed in pairwise combinations of 4 bacterial species relative to monoculture, where the addition of external nutrients led to increased competition, relative to the mutually beneficial interactions seen in a harsher environment (23).

The way in which species interact with one another not only depends on the resource environment but also on whether these originate from the same (sympatric) or a different (allopatric) environment. Sympatric bacteria are more likely to overlap in resource use and niche space (i.e. due to ecological filtering (24)) and are often more closely related, which tends to favour competitive interactions (13, 25–28). The higher relatedness of sympatric bacteria (29), could however lead to more positive interactions as a result of complementary cell surface adhesion and quorum sensing communication, although the reverse has also been found (30).

Laboratory growth media vary widely in resource complexity, from minimal media containing a single carbon source (e.g., M9) to rich, undefined media such as tryptic soy broth. These differences in resource composition are likely to favour different interaction outcomes by shaping resource use and niche overlap among competing strains. Additionally, pairwise competition experiments are often conducted in such laboratory media, designed to maximize bacterial cell densities. Under these conditions, selection for rapid growth—combined with ecological filtering for bacteria that share similar physiological traits (31, 32) —is likely to bias observed interactions toward more negative outcomes.

Crucially, laboratory media is likely to not be representative of a natural environment and may not allow the quantification of interactions which naturally occur. Therefore, systematic investigations of how bacterial interactions vary across different types of laboratory media, and relative to more natural conditions - whilst considering metabolic similarity driven by spatial co-occurrence - are critical.

Here, we tested whether interactions between bacterial pairs change between a semi-natural (soil wash) and two artificial growth media (Tryptic soy broth and minimal media), that differ in complexity. Furthermore, we tested the importance of geographic distance in driving the nature of species interactions.

We hypothesised that a rich and complex laboratory medium (Tryptic soy broth: TSB) would lead to high growth rates in monoculture and that this may lead to antagonistic interactions in coculture due to the ability to invest into growth and interference mechanisms (15), and could result in the loss of one competing isolate. Considering the more simplified resource of the minimal media, it was predicted that pairs may alleviate competition to synthesise the limiting resource (9, 10, 33) and result in less negative interactions relative to those seen in TSB. Finally, we predicted that soil wash would be an intermediate between the other two media and would allow the quantification of interactions more representative of those occurring in the natural environment. We predicted that sympatric bacteria (isolated from same soil sample) would experience more resource overlap and interactions would be largely competitive when compared to allopatric bacteria – particularly in the soil wash media, where the resource profile will match that of the species’ natural environment (34).

## Materials and Methods

### Bacterial isolation

Bacteria were sampled from four different sites, all contained within the gardens of the Penryn Campus, Penryn, Cornwall (50.170, 5.125). At each site, five soil samples were taken by fully inserting a sterile pipette tip (40mm) into the soil, after which tips were individually stored in 15 mL of minimal salts buffer. Soil suspensions were then plated onto King’s Medium B (KB) agar containing Nystatin (20 mg/L) to hamper fungal growth and incubated for 24 hours at 28°C. For each of the 20 plated soil samples, five morphologically dissimilar colonies were picked, yielding a total 100 bacterial isolates. Each isolate was regrown for 24 hours in TSB before being frozen in glycerol at a final concentration of 25% at -70°C.

All 100 isolates were tested for antibiotic resistance by selective plating onto KB agar supplemented with Gentamicin (20µg/L;10µg/L) and Tetracycline (10µg/L). To distinguish paired isolates that were visually similar, clones were paired such that one was antibiotic resistant and one susceptible. This allowed for stamp plating of coculture plates onto selective agar, and quantification of colony numbers of each isolate in pairwise assays. We note that selective plating was only used for three pairs as isolates within a pair were morphologically distinct on KB agar.

18 pairs of isolates were selected based on their antibiotic sensitivity (ensuring one resistant and one susceptible), with 6 pairs each taken from general, local and different environments (**Figure 1**). These three different localities are classed based on geographic proximity of pairs: ‘Local’ denotes isolates picked from the same plated community, originating from a single pipette tip; ‘General’ denotes isolates picked from different plated communities (different pipette tips), but which originated from the same site and ‘Different’ denotes isolates from different plated communities and sites.

**Figure 1:**
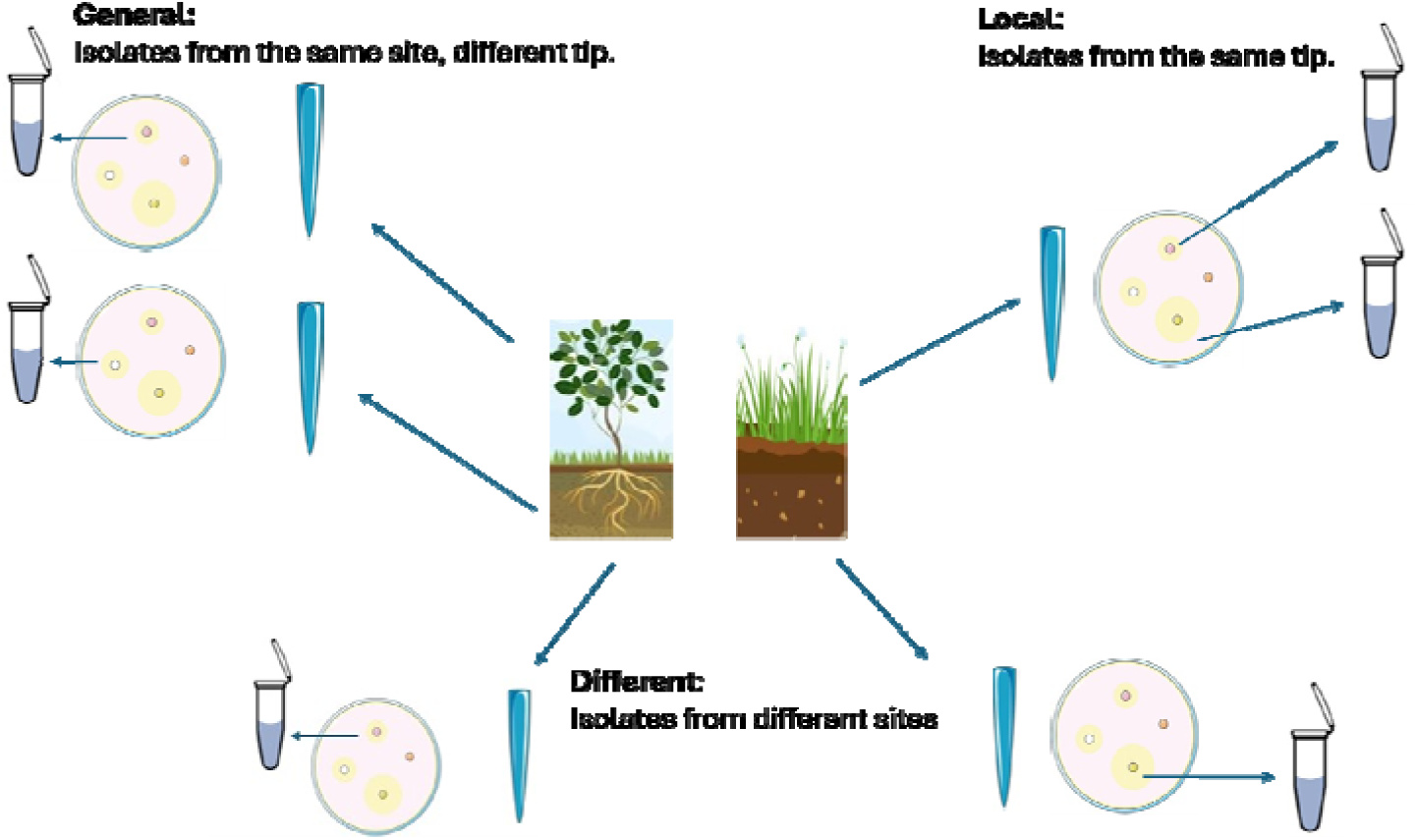
Bacterial isolation method showing how the three locality categories used relate to geographic proximity. Highest spatial overlap is the **local** treatment, where isolates were picked from the same plated community, originating from the same tip. **General** locality are isolates which are picked from different plated communities from different tips, but which were taken at the same site. The treatment **different** has the least spatial overlap, with isolates picked from different plated communities originating from different sites.

Values missing from subsequent plots were due to insufficient growth in either plated inoculum densities or following 48 hours of growth in media (isolates 21,22 and 35 removed from the study and 1 rep missing in isolates 16,33 and 34; **SI**).

### Experimental Procedure

To quantify how the resource environment affects bacterial growth and species interactions, we used three different media types: (1)TSB (Sigma-Aldrich), (2) minimal media supplemented with glucose (final volume 1 L: 20mL glucose solution, 2mL MgSO_4_ (1M), 0.1mL CaCL_2_ solution (1M), 100mL 10*M9 salts (**SI**), 878mL dH20) and (3) soil wash. The latter was made by collecting 200g of soil from the same general area the bacteria were isolated from and suspending this in 1L of water for 24 hours, after which the soil wash was filtered to remove larger particles and autoclaved twice with a 48-hour gap between both runs. It is important to note that the same soil wash was used for all replicates, and the collected soil used in the wash was not from any of the sampling locations but taken from a distinct soil environment within the same location.

To set up the experiment, single colonies of all selected isolates were grown statically in 6mL of TSB for 24 hours. To minimise resource transfer, 1mL of each culture was centrifuged for three minutes at 14000 rpm, after which we removed the supernatant, and resuspended the pellet into 1mL M9 buffer to remove access resources. These washed cultures were then used to inoculate the experimental media, and frozen in 50% glycerol at -70°C to determine inoculum densities. For monocultures, 60µL of resuspended culture was added to 6mL of relevant media in glass vials, for cocultures 30 µL of each member was added, keeping total inoculation density constant. The coculture treatment was repeated three times for each pair per medium. These were incubated statically at 28°C for two days after which all vials were vortexed and glycerol freezer stocks made. All freezer stocks were plated onto KB agar and colony counts were performed.

### Colony PCR and species identification

To understand the phylogenetic relationship of competing pairs, the 16S rRNA gene of each of the 26 isolates was sequenced, using 27F and 1492R primers and Sanger sequencing (Source Bioscience).

### Data Analyses

For all analyses R Version 4.3.1 was used. Model behaviour was tested using the ‘DHARMa’ package (35). Post-Hoc analyses were carried out using the ‘emmeans’ package (36), with p-values adjusted to correct for multiple testing, and all plots were made using ‘ggplot2’ (37), additional packages used are cited below. To determine the significance of variables, models were simplified by sequentially removing variables and compared via F, χ*^2^* or likelihood ratio tests (38), where appropriate.

To assess how the different environments affected isolate growth, we carried out a linear model, testing differences in monoculture density change in response to media type.

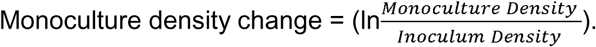

Where ‘Inoculum density’ is the inoculated colony counts of each isolate and ‘Monoculture density’ is the final colony counts following 48 hours growth in each media type.

For co-cultures, we quantified species interaction signs by calculating the Relative Intensity Index (RII) using mono- and co-culture colony counts for each isolate following 48 hrs of growth:

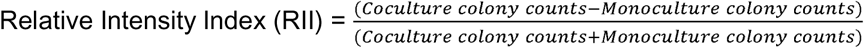

This proxy reduces the influence of extreme outliers arising in pairs where single isolates were outcompeted (39), with interaction indices for isolates bound between -1 and 1, such that positive values represent an increase in coculture relative to monoculture density and negative values reflect a decrease.

To test the effect of media type and locality on RII, a linear mixed effects model (Lmer) was fitted using the ‘lme4’ package (40), with a random intercept fitted for each clone and each pair to account for multiple observations. We tested for the main effects of media, locality and their two-way interaction – however this interaction did not significantly improve model behaviour and was subsequently removed from the final model.

To determine whether individual growth explains performance in pairwise co-culture, we carried out a linear model, testing the effect of media and monoculture growth, as well as their two-way interaction, on average RII, where Average RII = (RII_rep1_ + RII_rep2_ + RII_rep3_/3). A random intercept for ‘pairs’ was included, however the variance explained approach zero and its removal greatly improved model behaviour. The interaction fitted between media and monoculture growth did not significantly improve model behaviour and was subsequently removed from the final model.

Individual RII scores of paired isolates were then used to class interactions for each pair into mutually beneficial (+/+), exploitative (+/-) or competitive (-/-). For example, if isolate 1 RII = 0.1 and isolate 2 RII = -0.2, the pairwise interaction is classed as exploitative (+/-), with isolate 1 growing better in coculture relative to monoculture and the reverse being true for isolate 2. This was carried out for each individual replicate.

The likelihood of isolates having either mutually beneficial or exploitative interactions was assessed through a multinomial logistic regression, using the ‘nnet’ package (41), with the competitive interactions as base category, as this was the most frequent interaction seen across all isolates. Plotting and confidence intervals were carried out using ‘ggeffects’ (42).

Sanger sequences were analysed, using sangeranalyseR (1.20.0), where forward and reverse reads were trimmed, processed and aligned (43). Isolate 4 was unable to be sequenced, resulting in 35 sequenced isolates. For these 35 isolates, a pairwise distance matrix was created using the dist.dna function in the ‘ape’ package, and a phylogenetic tree constructed (44). Taxonomies shown on the phylogenetic tree were assigned using dada2, utilising the RDP Naïve Baysesian Classifier (45), taxonomies were unable to be assigned for 5 isolates. Chromatograms for each reaction and overall quality checks are available in the **SI**.

## Results

### Monoculture growth as a function of resource environment

To understand how the resource environment affected individual growth, we calculated the relative density change of each isolate following 48 hrs of incubation (Figure 2). Growth varied significantly across environments (linear model (lm), F= 64.557, P<0.001, Pairwise contrasts p<0.001 for all), with isolates growing fastest in TSB (**estimated mean** [95% confidence intervals], **3.47**, [2.84,4.10]). The minimal media decreased densities relative to the inoculum (-**1.69**, [-2.32,-1.05]) and the soil wash on average kept densities similar to the inoculum (**0.57**, [-0.05,1.19]).

**Figure 2:**
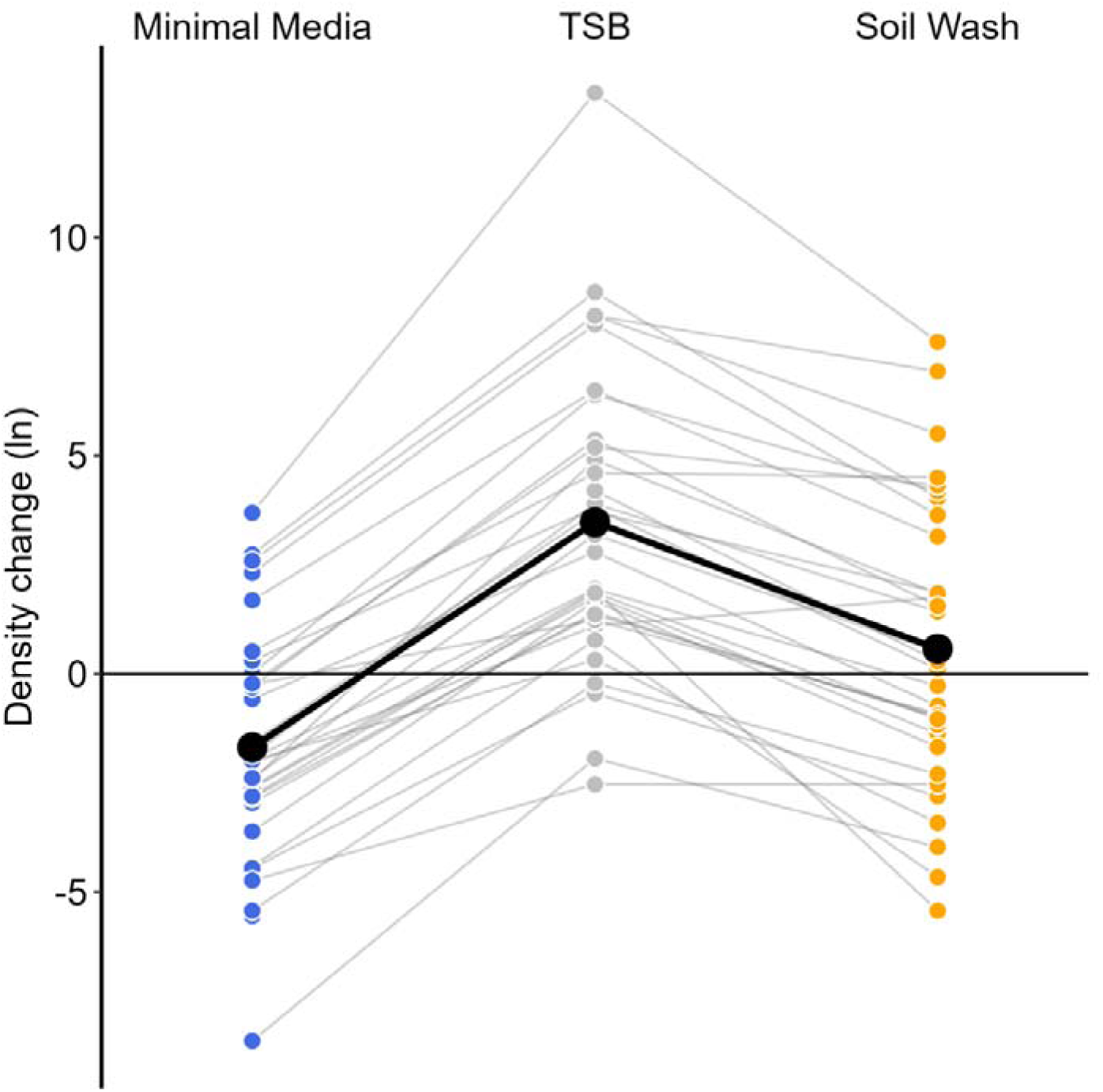
Plot showing individual growth (n = 36 isolates) as a function of resource environment. Growth was expressed as 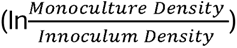. Grey lines link performance of individual isolates across environments, with the black line showing the mean growth. Blue dots show growth in minimal media, orange in soil wash and grey in Tryptic Soy Broth (TSB). Values above zero indicate isolates reached higher densities following 2 days incubation compared to that of the inoculum.

### Interactions vary across media types, independent of geographic proximity

Overall, negative interactions dominated across all treatments, with 195 negative interactions observed against 100 positive ones (i.e. uni-directional RIIs of individual isolates). Despite the prevalence of negative interactions, and within pair variation (which accounted for 25% of variation within the model: **S1**), resource environment significantly affected the nature of species interaction (Lmer on RII, media main effect, χ^2^=11.065, p=0.004, **Figure 3, S4**).

**Figure 3:**
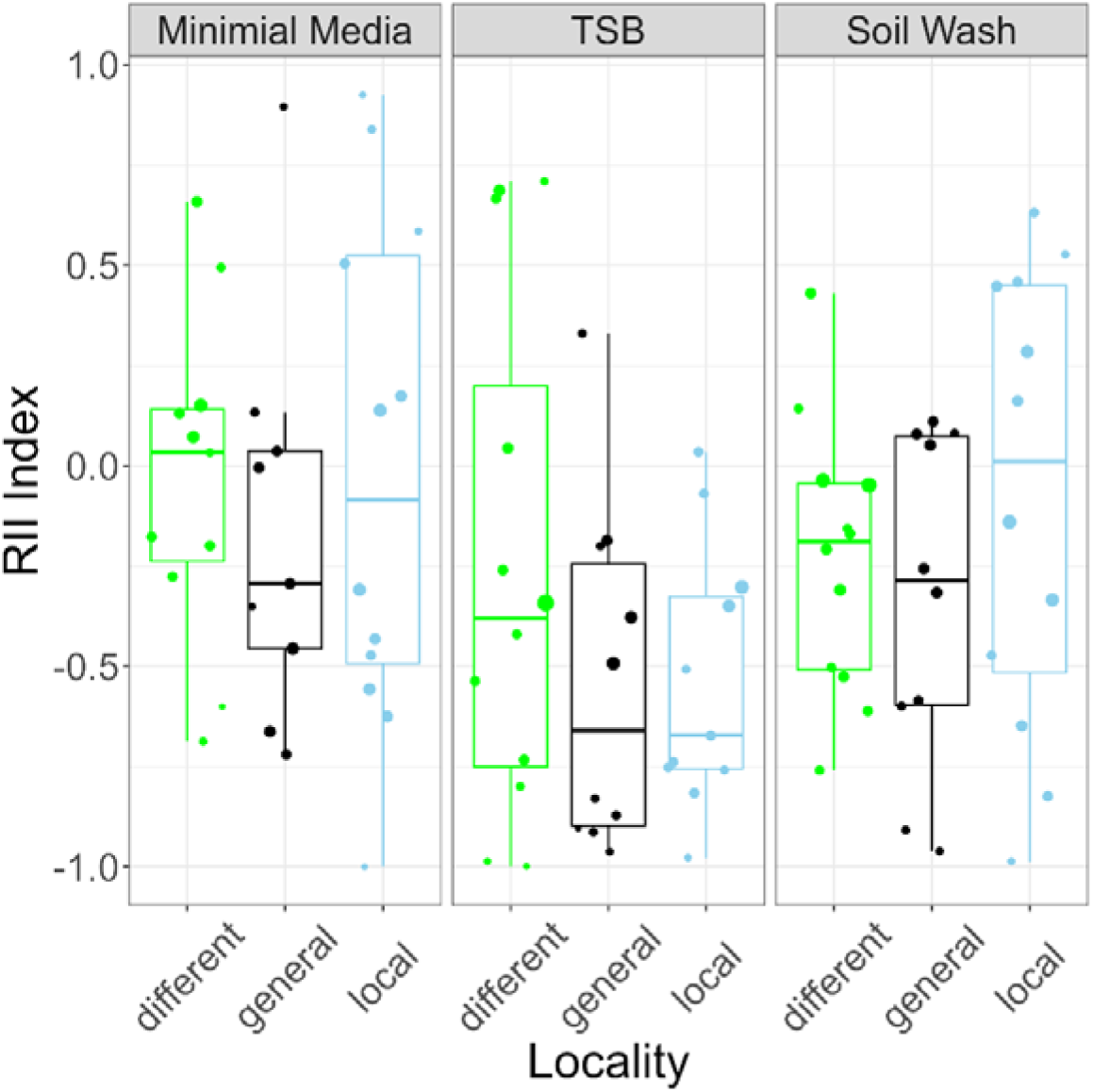
Boxplot showing the RII index of individual isolates as a function of resource environment (minimal media, TSB, Soil Wash) and geographic proximity (local, general, different) averaged across triplicates (n = 6 pairs and 12 isolates per unique treatment combination). Size of the data points depicts standard deviation around mean per isolate.

Isolates grown in TSB had the most negative interaction sign (**estimated mean**, [confidence intervals], **-0.44**, [-0.60, -0.27]), whilst those grown in minimal media were the least negative, with interactions on average being neutral (-**0.07** [-0.24, 0.09]). These resource environments differed most in terms of their resource abundance and complexity, which was reflected in a greater difference in RII index (post-hoc comparison: t=3.383, p=0.005, **S2**). This is in accordance with our predictions of more restricted resources favouring more positive interactions, and complex media allowing clones to invest in interference mechanisms and/or higher densities, thereby increasing competition. We therefore tested whether isolates with a greater propensity for growth in monoculture showed more negative RII indices in pairwise co-culture, and found this indeed to be the case (linear model, density change main effect, **estimated effect** [Standard error], **-0.056** [0.013], F_1,94_=19.794, P<0.001, **S3**) independent of the resource environment (resource x density effect, F = 0.5868, P = 0.558).

As predicted, interactions in soil wash tended to be less negative relative to TSB (**-0.20** [-0.37, - 0.04], soil wash-TSB post-hoc contrast: t=-2.184, p=0.088), and were not as positive as those seen in minimal media, though this difference was not significant (soil wash-minimal media post-hoc contrast, t=1.231, p=0.443).

We predicted sympatric isolates to interact more competitively than allopatric isolates due to similarities in resource use resulting from phylogenetic similarities. However, geographic proximity did not significantly affect bacterial interactions (Lmer on RII index: effect of media x locality interaction: χ^2^=3.827, p=0.430, locality main effect, (χ^2^=0.245 p=0.363)). Subsequent phylogenetic analysis showed that this was most likely due to most competing isolates within a pair belonging to different genera – even in pairs isolated in the same community (**Figure 4**) – such that phylogenetic diversity was similar across localities.

**Figure 4.**
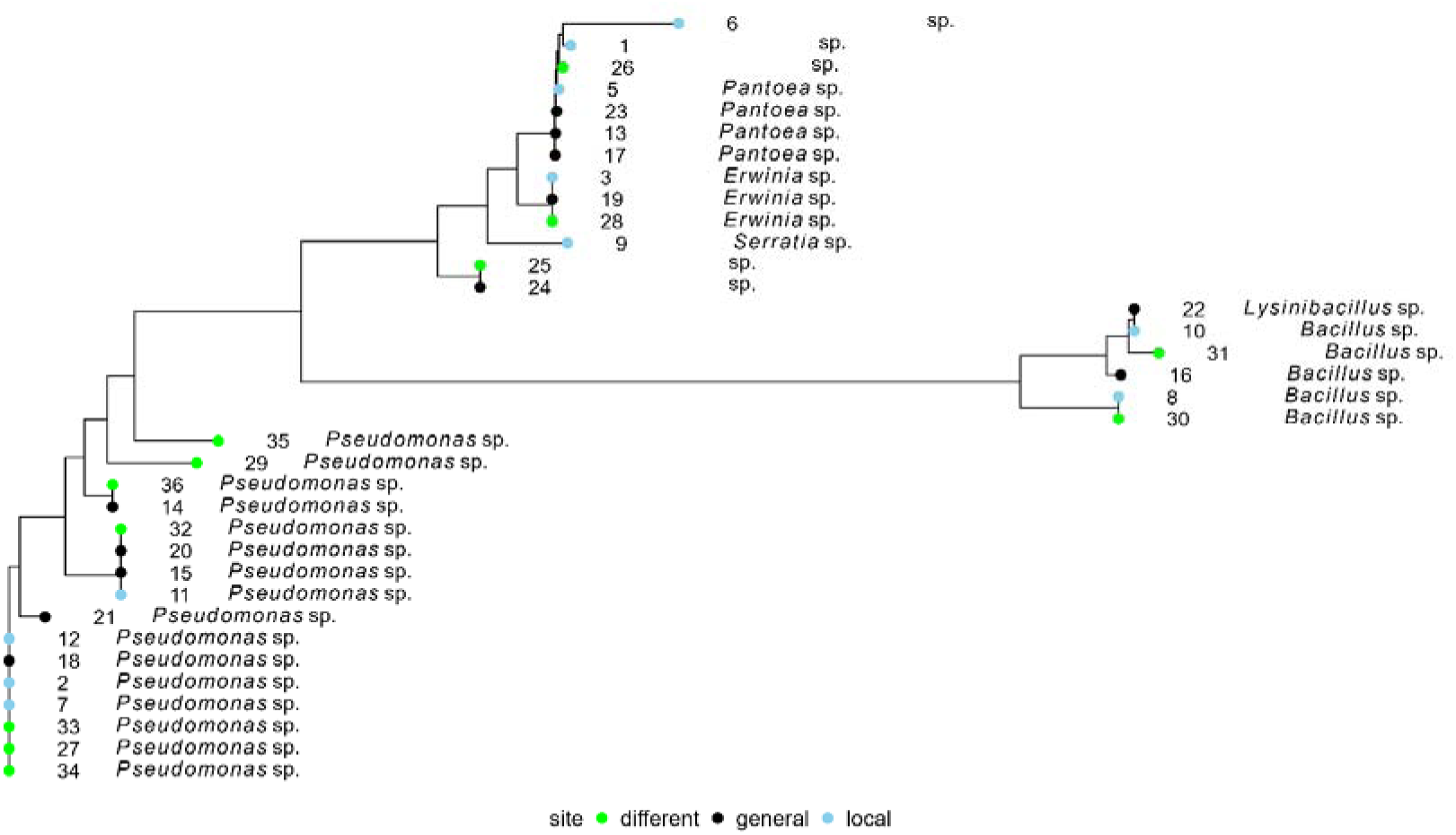
Phylogenetic tree showing genus level identification via 16S Sanger sequencing (see Methods) for each isolate. Pairs are made up in a stepwise fashion (e.g. pair 1: isolates 1 and 2; pair 2: isolates 3 and 4 etc). Some isolates could not be identified at the genus level, and for these, distances were calculated by aligning sequences. Isolate 4 is missing due to low quality reads.

The absence of a phylogenetic signal in response to location may reflect our sampling approach, which had limited ecological differentiation among sites. This may have promoted phylogenetic clustering through habitat filtering, thereby obscuring location-specific patterns.

### Reciprocal interactions depend on resource environment with exploitative interactions dominating in soil wash

We next classified reciprocal interactions by pairing the RII index of paired isolates as: cooperative (+/+), exploitative (+/-) or competitive (-/-) (see **methods**). Resource environment significantly affected the likelihood of different interactions (Likelihood ratio test, LR=12.411, p=0.015) Competitive interactions dominated in TSB (60% of total). However, there was a shift in reciprocal interactions in minimal media and soil wash towards mutualistic (+/+) interactions and exploitative interactions (+/-), respectively. (**Figure 5**).

**Figure 5:**
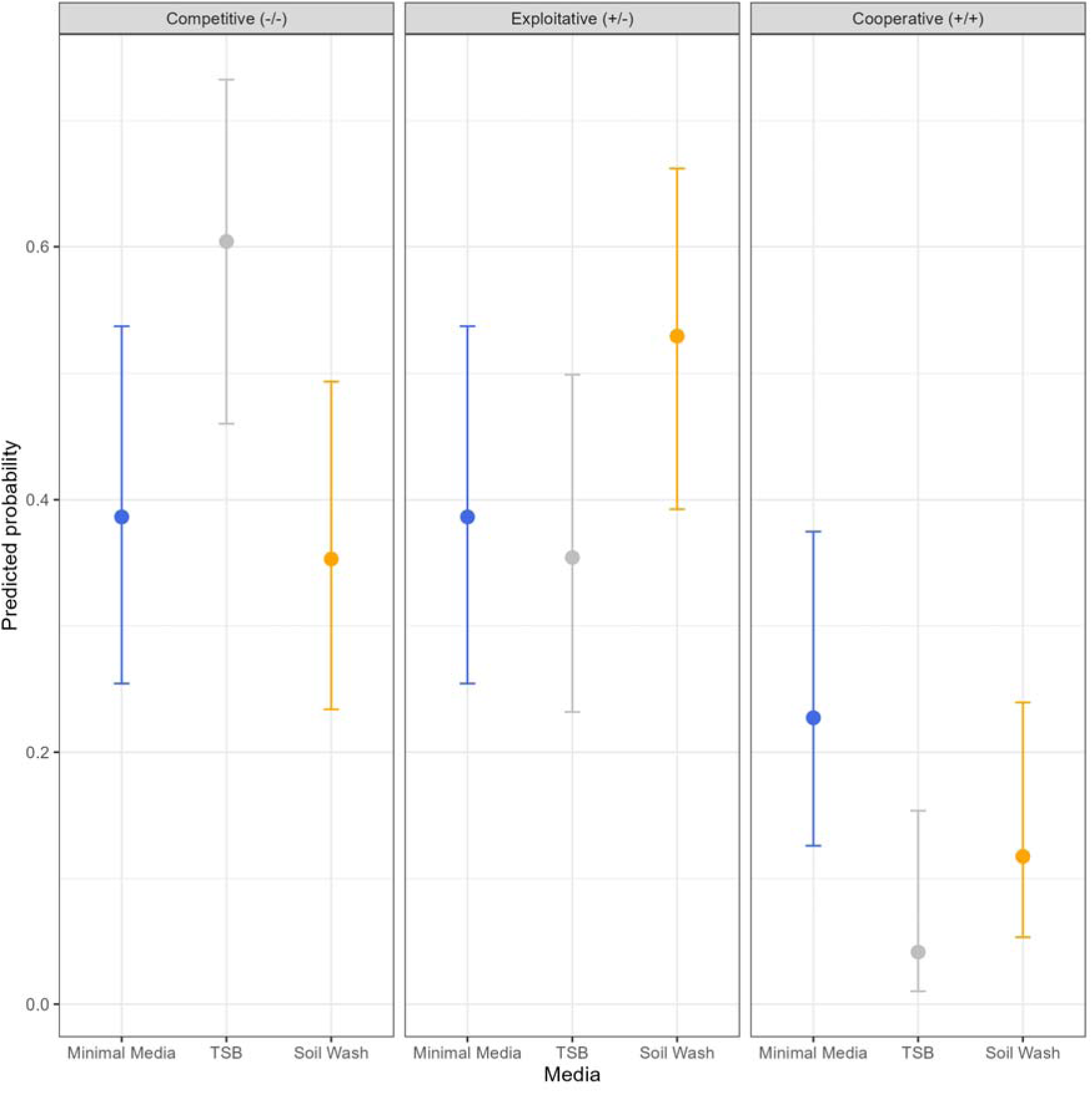
Plot showing the extracted probabilities predicted by the multinomial analysis with confidence intervals displayed above and below. The plot is facetted by interaction type and media shown on the x-axis.

To assess how species interaction strength was affected by resource environment, the absolute interaction index values (**Figure 6)** were used. This showed significant differences across environments in absolute isolate RII indices (LMER, media main effect, χ^2^=7.954, p=0.019, Figure 6). Interactions were strongest in TSB, where competitive (-/-) interactions prevailed, whilst interactions in soil wash were weakest overall.

**Figure 6:**
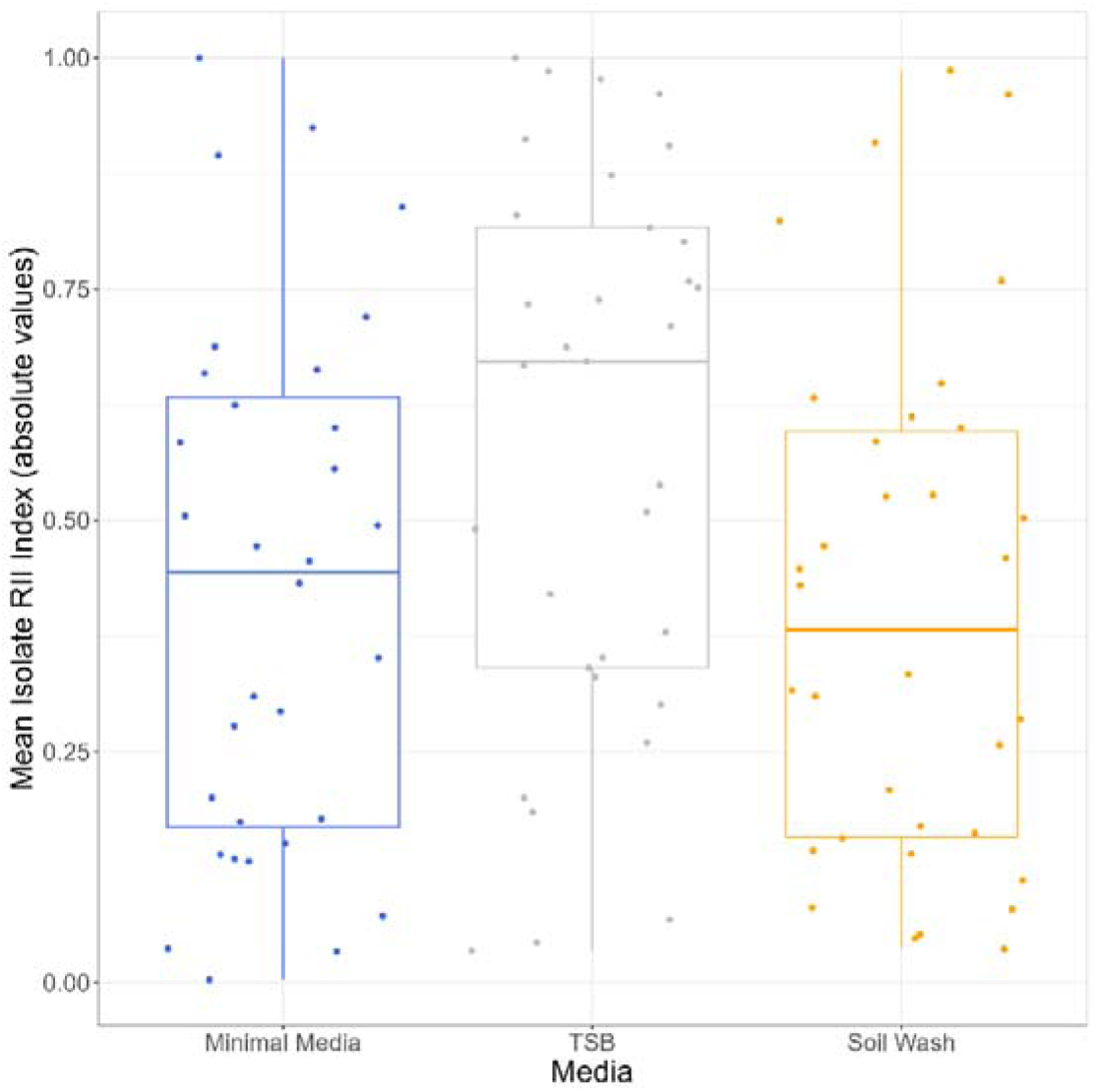
Boxplot showing the absolute values for the mean isolate RII index, given by the average isolate interaction value across the three experimental repetitions. All three levels of locality are combined so that the full data is plotted. Larger values represent stronger interactions.

## Discussion

We here tested the effect of environment type and geographic proximity on the nature of species interactions using a panel of environmental bacterial taxa isolated from different soil localities. Although we found competitive interactions to dominate, environments significantly affected the nature of species interactions: The most rich and complex environment (TSB) caused interactions to be significantly stronger and more negative, the few mutualistic interactions observed occurred primarily in the minimal medium, and the soil wash led to primarily exploitative and weaker interactions. In contrast to our expectations, interactions were independent of whether paired isolates originated from the same or different community/environment, which was likely due a lack of phylogenetic signal of locality, with phylogenetic diversity of isolates being equal across our locality treatments.

The lack of ecological differentiation among the sampled sites likely led to phylogenetic clustering because of habitat filtering and the reduced overall diversity seen in the sequenced isolates, with the majority being identified as *Pseudomonas sp.* Though isolates were not paired based on morphology, some sampling bias may have led to phylogenetically similar species as a result of picking colonies which looked morphologically distinct. In addition, the plating of soil communities on King’s Broth (KB) agar plates will have imposed some selective bias on all picked isolates.

Consistent with our predictions (11–13), competitive interactions dominated in TSB while the highest occurrence of mutualistic interactions occurred in minimal media. This is likely because the growth of some isolates was significantly constrained in minimal medium, leading to isolates engaging in cross-feeding behaviours when grown in coculture, utilising metabolic waste products of others (16). More generally, we found that an increase in individual growth was associated with more negative RII values (**S3**). In contrast with previous work (15), our findings suggest that increased growth, rather than interference investment, was the main driver underpinning competitive interactions in our study - with isolate specific trade-offs between investment in monoculture growth and survival in coculture potentially driving the trend seen, similar to that seen in cocultures of *S.marcescens* and *N.capsulatum* (21). These findings highlight that species interactions are highly context dependent, and that resource richness and diversity should be considered when studying species interactions in bacterial communities.

The soil wash was used to best reflect the isolates’ natural conditions. Consistent with previous work on natural isolates the highest proportion of exploitative interactions (+/-) occurred in this environment (46). Cross-feeding interactions in diverse soil communities are likely to consist of numerous uni-directional relationships, in which individuals attain higher densities through the utilization of by-product metabolites produced by neighbouring taxa. In addition, the type of cross-feeding interactions occurring in diverse soil communities are likely multiple uni-directional relationships, whereby individuals may reach higher densities because of by-product metabolite production from another individual (16). The exploitative interactions observed in soil wash communities may therefore arise from such cross-feeding dynamics, where accidental overproduction or metabolite leakage generates an asymmetric interaction in which one individual benefits from metabolic waste and potentially outcompetes the producing strain. The fact exploitative interactions in soil wash media were the weakest interactions overall is consistent with theory that exploitative interactions may increase stability in bacterial communities by reducing dependency between cooperators, and that weak interactions generally are more likely in natural communities (4, 5, 9)

Mutualistic interactions are predicted to be more common between isolates from different localities as inter-site resource differences increase (13, 30, 46, 47). Surprisingly, geographic proximity did not affect either the variation in interaction sign or the likelihood of a certain interaction being present in our study. This can in part be explained by the observed lack in phylogenetic-geographic relationship – there were few closely related pairs in our ‘local isolation’ category (and vice versa with a lack of distantly related pairs in pairs from different sites). Therefore, the similar phylogenetic diversity identified within and between sites likely masked any location effects, and although the sampled scale may ignore some natural complexity, small spatial differences have previously been shown to shape bacteria-phage local adaptation in soil environments (48).

## Conclusion

This study looked at the effect of three different environments of different resource complexity and richness on bacterial pairwise interactions across a panel of diverse soil bacteria. In addition, bacteria were isolated from different sites to test if geographic proximity affected interactions due to resource use overlap. We found that interactions were negative overall and varied across resources environments, but not with isolate locality.

Overall, this study demonstrated how a rich and complex laboratory medium leads to high growth rates and strong negative interactions, whilst showing how more simple resources lead to weaker interactions and in cases can lead to cooperation, though rare. Notably, the medium designed to mimic soil conditions was characterized by weak, predominantly exploitative interactions.

## Data availability

All data and code used in the study are available at (https://tinyurl.com/mr43x5ap)

