## supplementary info for "Sign and strength of pairwise interactions in natural isolates depend on environment type"

**Supplementary Information**

Four extra plots and recipe for 10*M9 salts, of which an aliquot was used to make up the minimal media used for the study.

S1 plot shows random effect plotted for each pair in media, with each pair plotted in each facet panel. S2 plot shows which contrasts are significant in response to using media as a predictor in modelling interaction indices. S3 shows how increases in monoculture leads to lower average RII (interaction index) overall. S4 shows the same data as figure 3, but with clones linked across the different media.

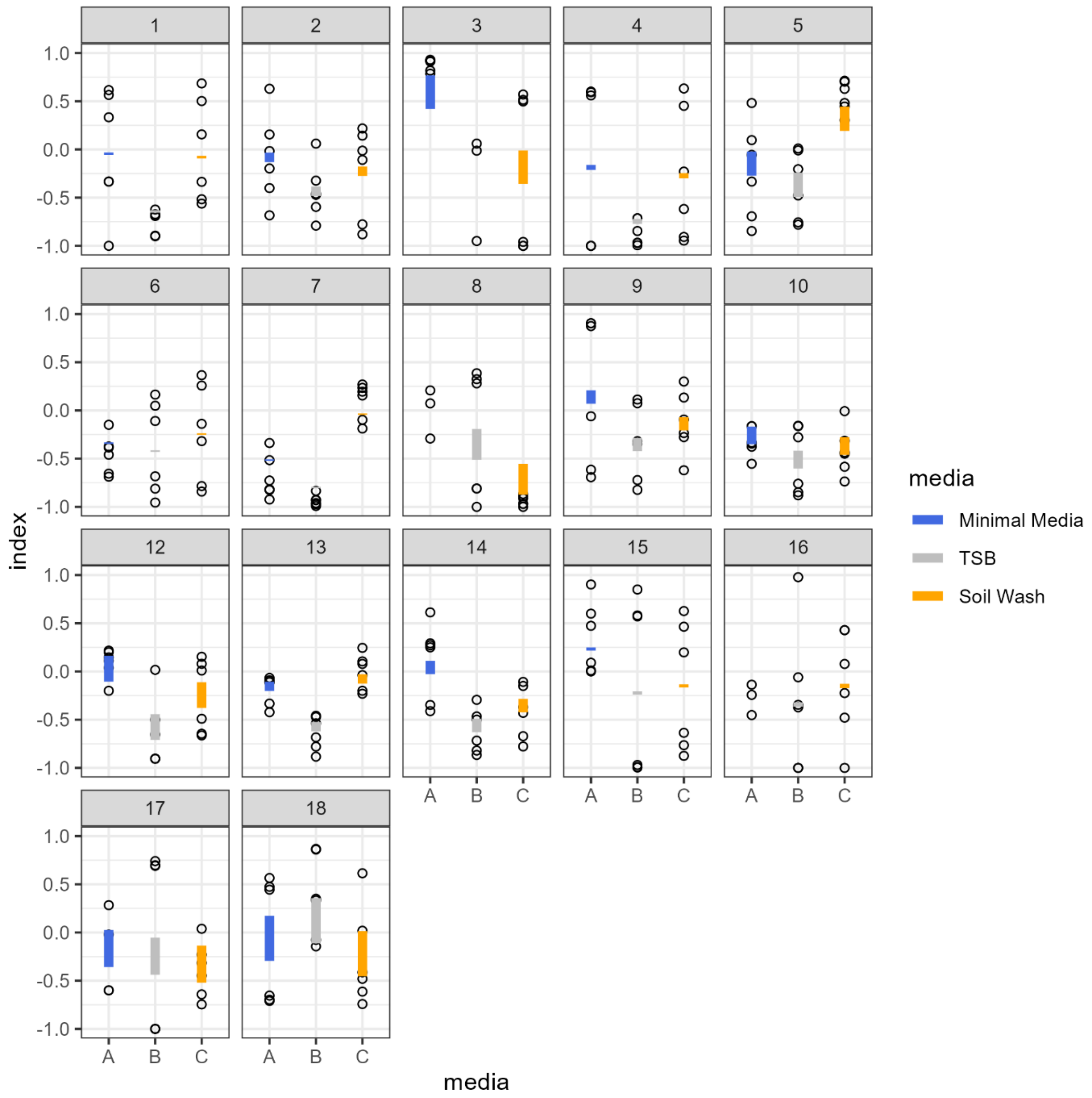


**S1**: Plot showing the interaction which was fit as a random effect in the final *LMER* model: Lmer (RII ~ media+locality +(1|pair.id:media) + (1|clone.id)). Black dots show raw data points (n=6 per pair) used in the model, whilst the coloured lines are conditional predictions layered and coloured by media (variation attributed to random effect in model). Each facet represents a bacterial pair.

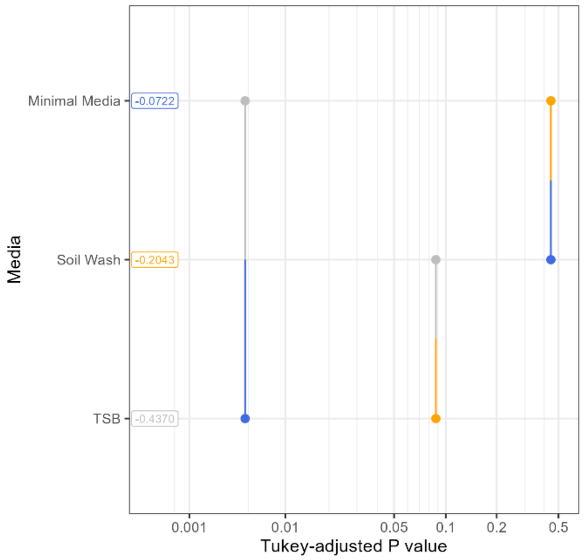


**S2:** Plot showing which contrasts are significant between media when using index as a response variable. Blue colour represents the minimal media; orange the soil wash and the grey is the TSB media.

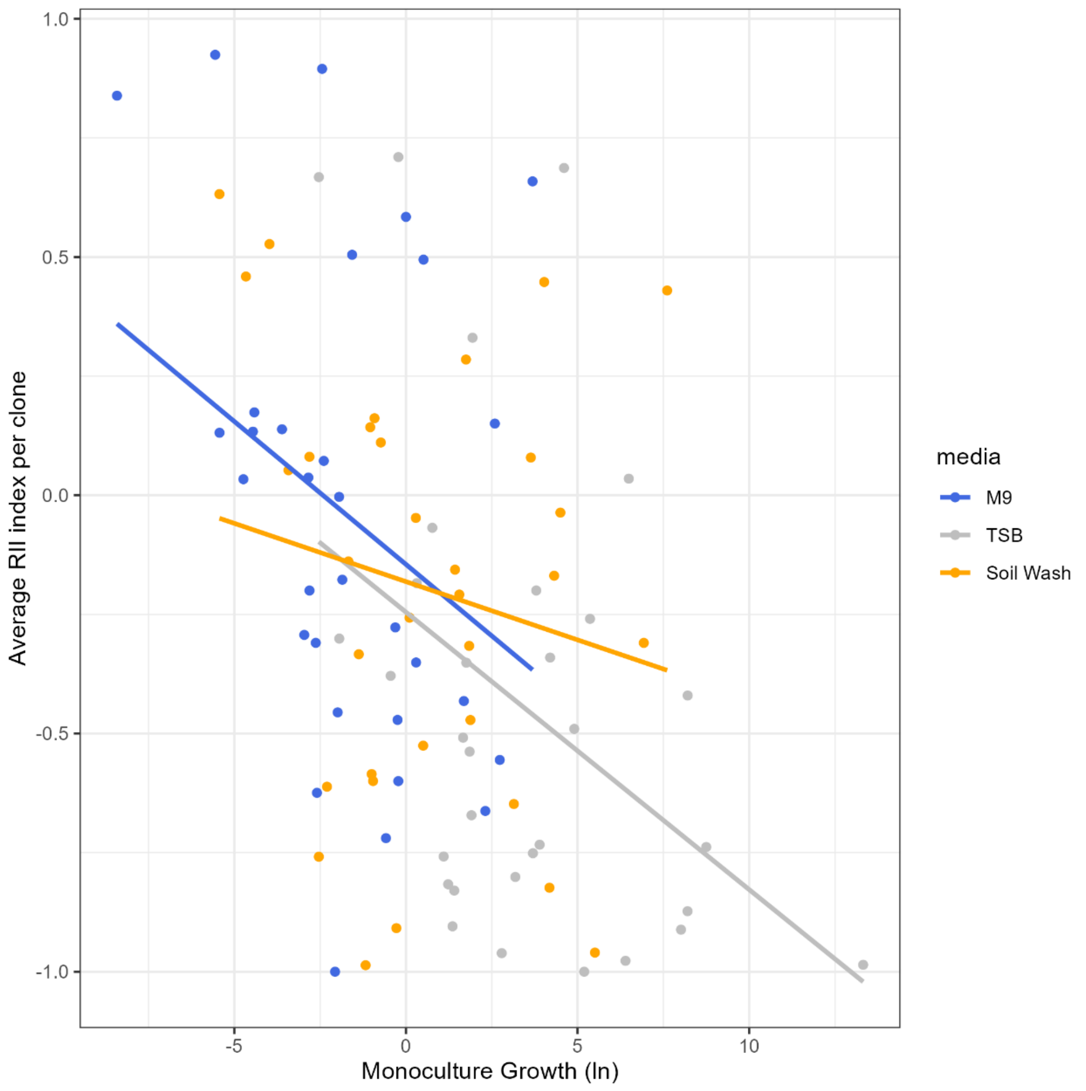


**S3**: Scatterplot showing how the average RII index per clone ($RII_{rep1} + RII_{rep2} + RII_{rep3}/3$) changes in response to increases in the density change seen in monoculture = ($\ln\frac{Monoculture\ Density}{Innoculum\ Density}$) for each isolate.

Increases in growth in monoculture led to more negative RII index across isolates (linear model, density change main effect, **estimated effect** [Standard error], **-0.056** [0.013], $F_{1,94}=19.794$, $P<0.001$) and this was consistent across media types (media x density change effect, F = 0.5868, P = 0.558).

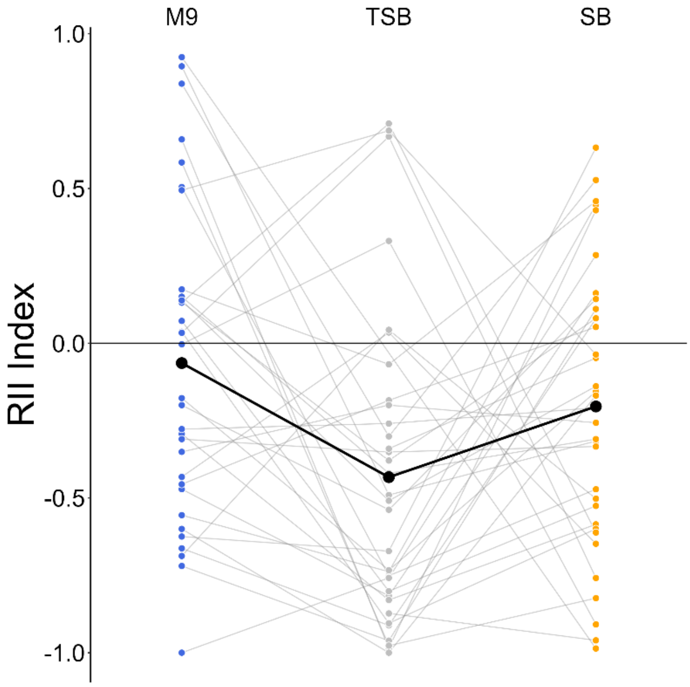


**S4** Plot showing the RII index for each isolate averaged across triplicates. Isolates are linked across media to aid visualization of media's effect on RII index. Mean values for each media are shown in black. Values close to zero indicate little change in density in coculture relative to monoculture, with more extreme positive and negative values representing large increase or decrease in density, respectively.

10*M9 Salts recipe used in minimal media: (7g $Na_2HPO_4.7H_2O$, 3g $KH_2PO_4$, 0.5g NaCl, 1g $NH_4Cl$ dissolved in 100mL $dH_2O$)
